# Dynamics of transposable element invasions with piRNA clusters

**DOI:** 10.1101/458059

**Authors:** Robert Kofler

## Abstract

In mammals and in invertebrates the proliferation of a newly invading transposable element (TE) is thought to be stopped by a random insertion of one member of the invading TE family into a piRNA cluster. This view is known as the trap model. Here we explore the dynamics of TE invasions under the trap model using large-scale computer simulations. We found that piRNA clusters confer a substantial benefit, effectively preventing extinction of host populations from an uncontrollable proliferation of deleterious TEs. We show that TE invasions under the trap model consists of three distinct phases: first the TE rapidly amplifies within the population, next TE proliferation is stopped by segregating cluster insertions and finally the TE is permanently inactivated by fixation of a cluster insertion. Suppression by segregating cluster insertions is unstable and bursts of TE activity may yet occur. The transpositon rate and the population size mostly influence the length of the phases but not the amount of TEs accumulating during an invasion. Solely the size of piRNA clusters was identified as a major factor influencing TE abundance. Investigating the impact of different cluster architectures we found that a single non-recombining cluster (e.g. the somatic cluster flamenco in Drosophila) is more efficient in stopping invasions than clusters distributed over several chromosomes (e.g germline cluster in Drosophila). With the somatic architecture fewer TEs accumulate during an invasion and fewer cluster insertions are required to stop the TE. The inefficiency of the germline architecture stems from recombination among cluster sites which makes it necessary that each diploid carries, on the average, four cluster insertions, such that most individuals will end up with at least one cluster insertion. Surprisingly we found that negative selection in a model with piRNA clusters can lead to a novel equilibrium state, where TE copy numbers remain stable despite only some individuals in a population carrying a cluster insertion. Finally when applying our approach to real data from *Drosophila melanogaster* we found that the trap model reasonably well accounts for the abundance of germline TEs but not of somatic TEs. The abundance of somatic TEs, such as gypsy, is much lower than expected.

## Introduction

Transposable elements (TEs) are short stretches of DNA that selfishly multiply within genomes, even when this activity has deleterious effects to the host [Orgel and Crick, 1980, Doolittle and Sapienza, 1980]. Deleterious effects may arise by three distinct mechanisms: i) TE insertions could directly disrupt genes or promotor regions, ii) ectopic recombination between insertions at different sites could lead to deleterious genomic rearrangements and iii) the products of TEs such as the transposase could be deleterious (eg. by generating double strand breaks) [Nuzhdin, 1999]. However, also several beneficial TE insertions, for example conferring resistance to insecticides, have been identified [Casacuberta and Gonzalez, 2013, Aminetzach et al., 2005]. On the overall, the fitness cost of TEs remain controversial. A recent review therefore argued that the null hypothesis for the fitness consequences of TE insertions should be the neutral model (i.e. a TE insertions have no or little effect on host fitness) [Arkhipova, 2018].

Due to the ability to proliferate within genomes TEs frequently invade novel populations and species [Kidwell, 1983, Kofler et al., 2015a, Peccoud et al., 2017]. There is ample evidence that an invasion of a TE may be triggered by horizontal transfer from a distant species [Kidwell, 1983, Kofler et al., 2015a, Rozhkov et al., 2013, Montchamp-Moreau, 1990]. It is likely that an invasion may also be triggered by processes that reactivate dormant TEs, such as environmental and genomic stresses, and by mutations within genes suppressing TE activity [Capy and Gibert, 2004, Prud’homme et al., 1995, Sarot et al., 2004, Kalmykova et al., 2005, McClintock, 1984, Wylie et al., 2016, Beauregard et al., 2008]. Irrespective of what triggered an invasio, an unchecked proliferation of TEs may drive host populations extinct [Brookfield and Badge, 1997], it is thus essential for organism to control the spread of TEs. It was long thought that the proliferation of TEs is counteracted at the population level by natural selection acting against deleterious TE insertions [Charlesworth and Charlesworth, 1983, Charlesworth and Langley, 1989, Barrón et al., 2014]. According to this “transposition-selection balance model”, TE copy numbers within a population are at an equilibrium between transposition events generating new insertions and negative selection removing insertions [Charlesworth and Charlesworth, 1983, Charlesworth and Langley, 1989, Barrón et al., 2014].

However, the discovery of the small-RNA based defence system profoundly changed our view on TE dynamics. It showed that the spread of TEs is not solely counteracted at the population level but actively combated by the host [Blumenstiel, 2011, Lee and Langley, 2010]. The host defence system relies on the so called piRNAs, small RNAs ranging in size from 23 to 29nt [Brennecke et al., 2007, Gunawardane et al., 2007]. piRNAs bind to PIWI-clade proteins and mediate the suppression of TEs at the transcriptional and at the post-transcriptional level [Sienski et al., 2012, Le Thomas et al., 2013, Brennecke et al., 2007, Gunawardane et al., 2007]. piRNAs are largely derived from discrete genomic loci that have been termed piRNA clusters [Brennecke et al., 2007, Malone et al., 2009]. These piRNA cluster are frequently found in the heterochromatin, close to the euchromatin boundary, and may make up a substantial fraction of genomes [Brennecke et al., 2007]. For example in *Drosophila melanogaster*, piRNA clusters constitute about 3.5% of the genome [Brennecke et al., 2007]. Several studies found that a single TE insertion in a piRNA cluster may be sufficient for repressing the activity of a TE [Josse et al., 2007, Zanni et al., 2013, Ronsseray et al., 1991]. Such observations gave rise to the “trap model”, which holds that an invading TE proliferates within a host until at least one copy jumps into a piRNA cluster (the trap), which triggers production of piRNAs that silence the invading TE [Bergman et al., 2006, Malone and Hannon, 2009, Zanni et al., 2013, Yamanaka et al., 2014, Goriaux et al., 2014].

Interestingly, TEs may employ different strategies to increase in copy numbers [Blumenstiel, 2011]. They may either be active directly in the germline or in the somatic tissue surrounding the germline. Somatic TEs usually require virus like particles to infect the germline [Song et al., 1997]. Notably these two different groups of TEs may be controlled by two different specialized piRNA pathways that rely on distinct sets of piRNA-clusters [Malone et al., 2009, Li et al., 2009]. These two sets of piRNA clusters may further have distinct architectures [Malone et al., 2009, Li et al., 2009]. In *D. melanogaster* somatic TEs are controlled by a single piRNA-cluster, flamenco, which is located in heterochromatic regions of the X-chromosome whereas germline TEs are controlled by several piRNA-clusters distributed over multiple chromosomes [Brennecke et al., 2007, Malone et al., 2009]. Additionally, TE insertions in flamenco are overwhelmingly in an antisense orientation while no such bias was found for insertions in germline clusters [Malone et al., 2009].

piRNAs and piRNA-clusters have been found in many different species such as flies, worms, mouse and humans [Lewis et al., 2018, Yamanaka et al., 2014, Aravin et al., 2007, Czech and Hannon, 2016]. It is therefore likely that the trap model holds for most invertebrates and mammals. Despite this wide applicability the dynamics of TE invasions under the trap model have been little explored [but see Kelleher et al., 2018, Li et al., 2009]. Therefore we performed large-scale simulations of TE invasions under the trap model using our novel simulator Invade (https://sourceforge.net/projects/invade/). We show that piRNA clusters are highly beneficial to host populations as they prevent extinction from an uncontrollable proliferation of deleterious TEs. We furthermore show that TE invasions have three distinct phases. We found that the size of piRNA clusters is the most important factor governing the amount of TEs accumulating during an invasion and that the somatic architecture is more efficient in stopping invasions than the germline architecture. Finally, using publicly available data from *D. melanogaster* we found that the trap model reasonably well accounts for the abundance of germline TEs but fails to explain the abundance of somatic TEs.

## 1 Results

The trap model holds that proliferation of an invading TE is stopped by a random insertion into a piRNA cluster (fig. 1A). In this work we performed large-scale simulations to gain a deeper understanding of the population dynamics of TE invasion under the trap model. We simulated 5 chromosomes with a size of 10Mbp, a recombination rate of 4cM/Mb and a piRNA cluster of size 300kb at the beginning of each chromosome (fig 1B). Thus, similarly as in Drosophila, the total size of the piRNA-cluster accounts for 3% of the genome [Brennecke et al., 2007]. Per default we used a population size of N = 1000. We launched TE invasions by randomly distributing 10 insertions in individuals of the starting population. This however raises a problem. Since the TE insertions in the starting population are initially segregating at a low frequency (1/2*N*) it is feasible that all TE insertions may be lost due to drift in some of the replicates [Le Rouzic and Capy, 2005]. The probability of loosing a TE insertion with frequency 1/2*N* is approximately *p* = 1 − 2*u* with *u* being the transposition rate [Le Rouzic and Capy, 2005]. Hence, the probability of successfully establishing a TE invasion is *p_e_* = 1 − (1 − 2*u*)^*n*^ with *n* being the number of insertions in the starting population. Using for example the different transposition rates *u* = 0.01, *u* = 0.1 and *u* = 1.0 we obtain probabilities of establishments of *p_e_* = 0.183, *p_e_* = 0.893 and *p_e_* = 1.0 respectively (*n* = 10). Our simulations agree with this expectation. Out of 1000 simulations the invasion got established in 168, 803 and 1000 replicates which is close to theoretical expectations (183, 893, and 1000, respectively; 500 generations were simulated). Since we are mainly interested in the dynamics of successful TE invasions we henceforth ignore TE invasion that failed to establish (unless mentioned otherwise).

**Figure 1:**
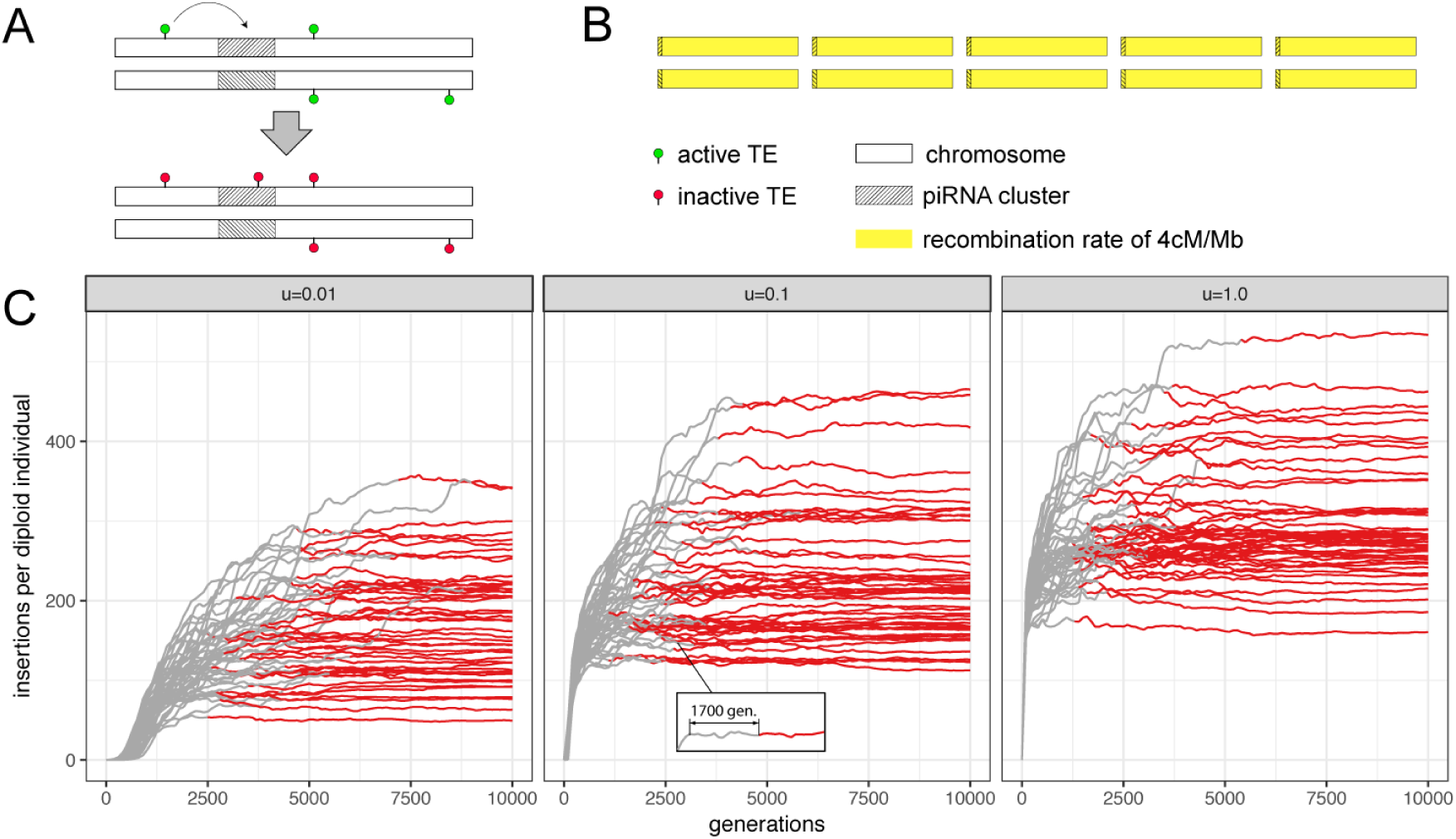
piRNA cluster may stop TE invasions. A) Under the trap model, an active TE (green) multiplies within the genome (rectangles indicate chromosomes of a diploid organism) until one copy jumps into a piRNA cluster (i.e. the trap, hatched area) whereupon all TEs, including those on homologous chromosomes, get inactivated in trans (red). B) We simulated 5 chromosomes of size 10Mb for a diploid organism. Each chromosome carried a piRNA cluster of size 300kb. A constant recombination rate of 4 cM/Mbp (yellow) was used. C) Abundance of TEs during an invasion. Populations of size *N* = 1000 and neutral TE insertions were simulated. We show 50 replicates for three different transposition rates (*u*: top panel). All populations eventually acquired a fixed cluster insertion (red line), which permanently inactivates the TE. Negative selection against TEs is thus not necessary to stop TE invasions under the trap model.

Classic population genetic models (transposition-selection balance), developed before the discovery of piRNAs, show that the proliferation of TEs can be contained by negative selection against TEs [Charlesworth and Charlesworth, 1983, Charlesworth and Langley, 1989]. We first tested the hypothesis that piRNA clusters are also capable of containing the spread of TEs. We simulated 100 TE invasions for 20.000 generations using three different transposition rates (*u* = 0.01, *u* = 0.1, *u* = 1.0; fig. 1C). Initially we simulated neutral TE insertions (i.e. TE insertions have no fitness costs to the host). Negative selection against TEs is treated later. Here, we define a TE invasion to be ‘‘stopped” once a cluster insertion gets fixed, as this permanently inactivates the TE. By generation 20.000 all replicates for each transposition rate acquired at least one fixed cluster insertion (trajectories for 50 replicates and 10.000 generations are shown in fig. 1C). This is in contrast to the results of Kelleher et al. [2018] who found that cluster insertions rarely got fixed. The discrepancy is likely due to the smaller number of generations used by Kelleher et al. [2018] (500 generations versus 20.000 in this work). We conclude that the piRNA clusters are able to stop TE invasions, even when transposition rates are extremely high and TE insertions are neutral (fig. 1C).

We noticed that in some replicates TE copy numbers stabilized for many hundred generations despite no cluster insertion being fixed (fig. 1C inlay). This suggests that the TE invasion may be contained by segregating cluster insertions as proposed previously [Kelleher et al., 2018, Kofler et al., 2018]. We therefore investigated the early stages of TE invasions in more detail. Interestingly, we found that TE copy numbers plateaued in all replicates although no cluster insertion got fixed (fig. 2A; *u* = 0.1). The average amount of novel TE insertions per generation and individual significantly decreased from 1.001 at generation 100 to 0.043 at generation 500 (Wilcoxon rank sum test; *W* = 9895, *p* < 2.2*e* − 16; fig 2B). This plateauing of the invasion was accompanied by an increase in the average amount of cluster insertions per individual, from 0.85 at generation 100 to 5.12 at generation 500 (Wilcoxon rank sum test; *W* = 0, *p* < 2.2*e* − 16; fig 2A). At early stages of the invasions all cluster insertions segregate at a low frequency, whereas high frequency insertions emerge at later stages (fig. 2C; supplementary fig. 2). Accordingly most cluster insertions were heterozygous at early stages of the invasions (fig. 2D). Our results thus support the view that TE invasions are initially stopped by segregating cluster insertions. However, at later stages fixed cluster insertions emerge (fig. 2C; generation 5000) which inactive the invading TE (fig. 2B; red dots indicate populations with fixed cluster insertions). Hence our results suggest that TE invasions under the trap model consist of three distinct phases. First TE copy numbers rapidly increase fairly unconstrained. We termed this the “rapid invasion” phase (fig. 2E, green). Second TE invasions are contained by segregating cluster insertions. Consistent with our previous work we term this the “shotgun phase”, to signify that cluster insertions are widely distributed over many distinct genomic sites (fig. 2E, yellow [Kofler et al., 2018]). Delimiting the exact onset of the shotgun phase is however a bit arbitrary. In this work we use the generation at which 99% of the individuals acquired at least one cluster insertion as the onset of shotgun phase. Invasion considerably slowed down at this stage (fig. 2A; dashed line). Third, fixation of a cluster insertions leads to a complete inactivation of the TE. Hence we termed this the ‘‘inactive” phase (fig. 2E red). We found that the number of insertion sites in the population increased sharply during the rapid invasion phase, decreased slowly during the shotgun phase and stabilized in the inactive phase (supplementary fig. 1). The site frequency spectrum of cluster and non-cluster insertions was identical during the invasion (ANOVA, effect of cluster vs. non-cluster category *p* = 0.97; supplementary fig. 2).

**Figure 2:**
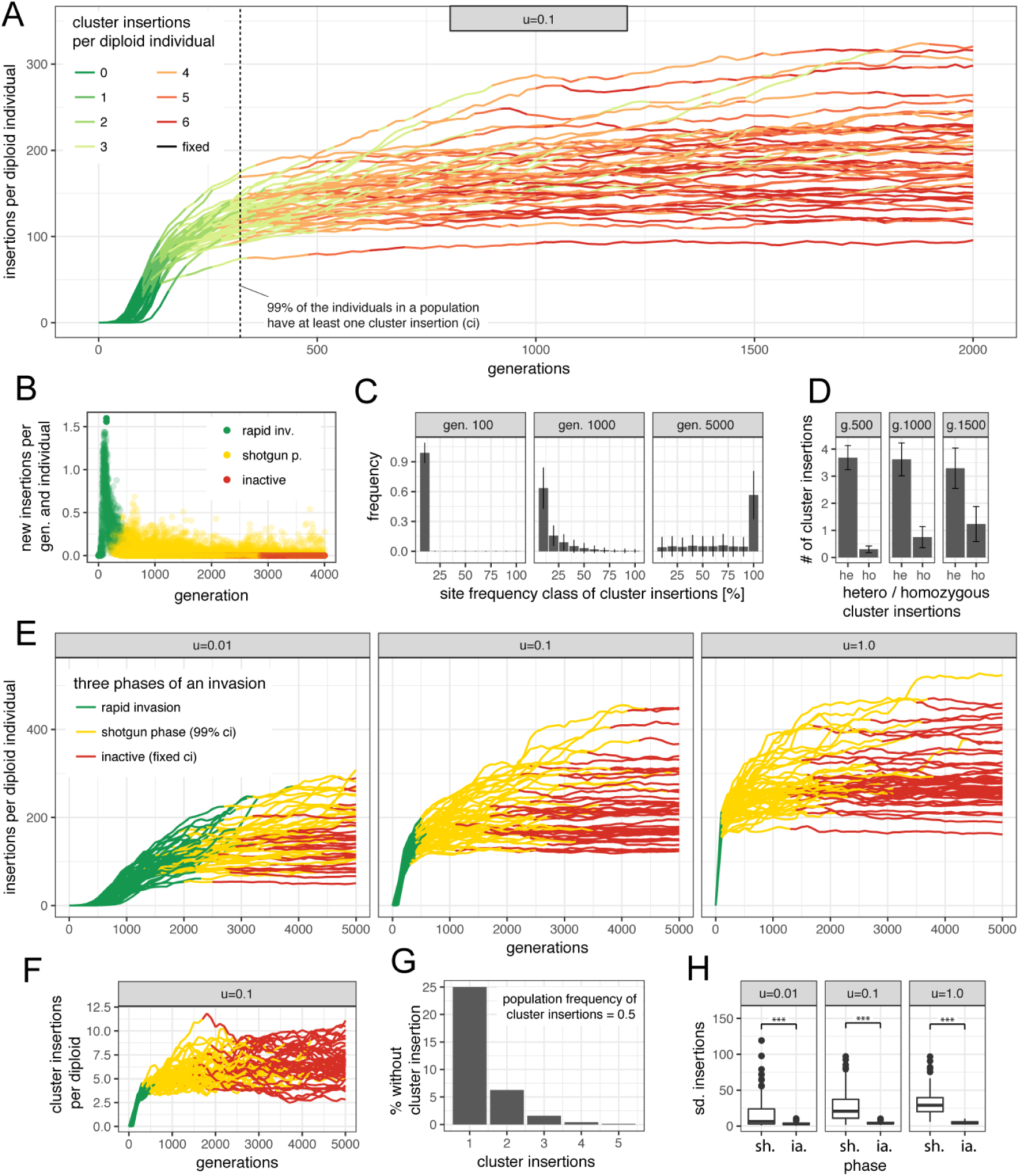
TE invasions consist of three distinct phases. A) Under the trap model TE invasions are initially stopped by multiple segregating cluster insertions. The invasion slows down as the number of cluster insertions per individual increases. Dashed line indicates the generation at which >99% of the individuals acquired at least one cluster insertion. No fixed cluster insertions were observed by generation 2000. B) Number of novel insertions per individual during TE invasions. TE activity ceases completely once a cluster insertion got fixed (red dots). C) Site frequency spectrum of cluster insertions during TE invasions. At early stages of an invasion (e.g. generations ≤1000) all cluster insertions segregate at low frequency. Error bars indicate standard deviation based on 100 replicates. D) Fraction of homo- (ho) and heterozygous (he) cluster insertions at different generations (g). E) The three phases of TE invasions for different transposition rates (*u*). Fifty replicates are shown. F) Number of cluster insertions for the three phases of TE invasions. G) Fraction of individuals without cluster insertions (i.e with an active TE), dependent on the number of segregating cluster insertions. H) Stability of phaišes measured in standard deviation (sd.) of TE copy numbers. The shotgun phase (sh.) is significantly less stable than the inactive phase (ia.; *** *p* < 0.01).

Interestingly we observed that at the onset of the shotgun phase each individual had on the average acquired 3.8 cluster insertions (e.g. with *u* = 0.1; fig. 2F), although a single insertion would have been sufficient to silence the TE. This can be explained by the fact that cluster insertions are segregating. Assuming a scenario where a single cluster insertion has a population frequency of 0.5. Due to Hardy-Weinberg equilibrium 25% of the individuals will not have a cluster insertion. Extending this example to two cluster insertions at distinct genomic sites, then of the 25% of individuals without cluster insertion at the first locus, another 25% will not have an insertion at the second locus. The TE will thus be active in the 6.25% of individuals without cluster insertion (fig. 2G). The fraction of individuals with an active TE can thus be computed as *f_a_* = Π_*i*_(1 − *p_i_*)^2^ where *p_i_* is the population frequency of the *i^th^* cluster insertion. On the average 3.8 cluster insertions per diploid are thus necessary to reduce the fraction of individuals with an active TE sufficiently such that TE copy numbers stagnate.

We noticed that in some replicates TE copy numbers increased abruptly during the shotgun phase (fig. 3E). To quantify the stability of the phases we computed the standard deviation of the TE abundance for each phase and replicate (fig. 3A right panel). We found that TE abundance during the shotgun phase is significantly less stable than during the inactive phase (Wilcoxon rank sum test; *p* < 4.1*e* − 08 for *u* =1, *u* = 0.1 and *u* = 0.01; fig. 3H). Our results thus show silencing of TE invasion by segregating cluster insertions is instable. Solely fixation of a cluster insertions results in permanent inactivation of the TE and thus in stable TE copy numbers.

Next we asked which factors influence the dynamics of TE invasion under the trap model. We evaluated the impact of the transposition rate (*u*), the genome size, the size of the piRNA clusters (in percent of the genome size), the population size (*N*) and the excision rate (*v*). To minimize the parameter space for the simulations we used default conditions (*u* = 0.1, genome size = 50*Mb*, cluster size=3%, *N* = 1000, *v* = 0%) and varied only the parameter of interest within these defaults (fig. 3; defaults are shown bold). We assessed the impact of these factors on the following key properties of invasions: the length of the phase, the TE abundance at the beginning of the phase, the abundance of cluster insertions at the beginning of the phase and the stability of the phase (quantified as standard deviation of the TE abundance per phase and replicate). We omitted meaningless or irrelevant data such as the length of the inactive phase (infinite) or the TE abundance at the beginning of the rapid invasion phase (10/2 * *N*) (fig. 3). We found that the transposition rate had a strong influence on the length of the rapid invasion phase but little influence on other properties, including the abundance of TE insertions (fig. 3)A; supplementary table 1). This is notable as the transposition rate is a major factor governing TE abundance under the transposition-selection balance model [Charlesworth and Charlesworth, 1983, Kofler et al., 2015b]. As expected the genome size had very little influence on the invasion dynamics (fig. 3)B; supplementary table 1). The reason why it had any influence at all may be that we ignored insertions into already occupied sites. Such double insertions are more likely to occur in smaller genomes and as a consequence fewer TEs will accumulate in smaller genomes. The size of the piRNA clusters had an enormous influence on the number of TEs accumulating during an invasion, where most TEs were found for small clusters (fig. 3)C; supplementary table 1). With small clusters many more insertions will be necessary until one copy randomly jumps into a piRNA cluster. This is also in agreement with previous works [Kelleher et al., 2018]. Interestingly the population size had a pronounced influence on the length of the shotgun phase, with larger populations having longer shotgun phases (fig. 3)D; supplementary table 1). Genetic drift is weak in large populations. Hence fixation of cluster insertions, which marks the end of the shotgun phase, will require more time than in small populations. Due to the longer duration of the unstable shotgun phase more TEs will accumulate in large populations than in small populations (fig. 3)D). Note that this result is in stark contrast to the classic transposition-selection balance model, where fewer TEs are expected to accumulate in large populations as the efficacy of negative selection against TEs is higher in large populations [Charlesworth and Charlesworth, 1983, Kofler et al., 2015b]. The excision rate only had a small influence on invasion dynamics (fig. 3)E; supplementary table 1). Also the recombination rate only had a weak influence on invasion dynamics (supplementary fig. 3; supplementary table 1). Surprisingly, we found that irrespective of the simulated scenario always about 4-6 cluster insertions per diploid where necessary to stop the invasions (fig. 3)). piRNA clusters should thus contain multiple insertions from silenced families.

**Figure 3:**
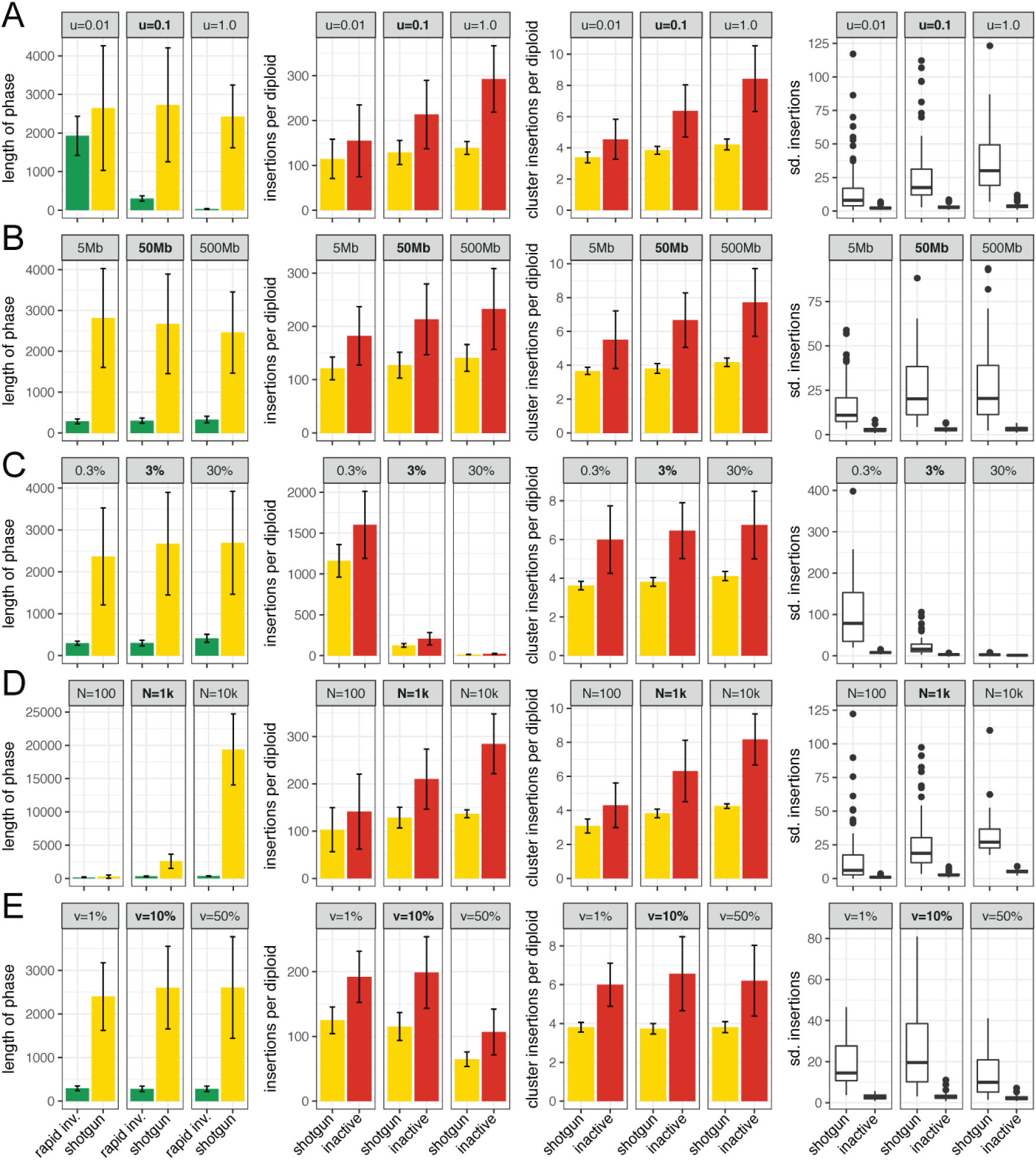
Influence of different factors on TE invasions. We studied the influence of the transposition rate (A), the genome size (B), the size of piRNA clusters, in percent of the genome (C), the population size (D) and the excision rate (E). For all simulations we kept default parameters (bold) and varied solely the factor of interest. We show the impact of the different factors on the length of the phase (in generations), the TE abundance per diploid individual at the start of the phase, the number of cluster insertions per diploid individual at the start of the phase and the stability of phase measured in standard deviation of the TE abundance (sd. insertions).

piRNA clusters, even within a given species, may have profoundly different architectures. For example in Drosophila two specialized piRNA pathways exist which rely on different sets of piRNA clusters [Li et al., 2009, Malone et al., 2009]. The somatic pathway mostly relies on a single cluster, i.e. flamenco, which is located in low recombining regions of the X-chromosome. The germline pathway, on the other hand, relies on several clusters (≈142) that are distributed over multiple chromosomes [Brennecke et al., 2007]. We hypothesized that this difference in architecture may have an impact on invasion dynamics. To test this we simulated 5 chromosomes with a size of 2Mb, a piRNA cluster size of 1Mb (i.e. 10% of the genome) and varied the number, the recombination rate and the genomic location of the clusters, while keeping the total size of piRNA clusters constant (fig. 4A). A single cluster in non-recombining regions resembles the somatic architecture (flamenco-model) and multiple clusters distributed over five chromosomes resembles the germline architecture (germline-model; fig. 4A). For each architecture we simulated 100 replicates. We found a pronounced difference of TE invasion dynamics between the flamenco and germline model (fig. 4B; supplementary table 2). While the length of the rapid invasion phase is significantly longer in the germline-model the length of the shotgun phase is significantly longer in the flamenco-model (fig. 4C; supplementary table 2). Notably, the number of TE insertions accumulating during an invasion is much lower in the flamenco-model than in the germline-model (fig. 4D; supplementary table 2). Also the number of cluster insertions necessary to stop an invasion is significantly lower with the flamenco-model (fig. 4E; supplementary table 2). Finally the stability of the shotgun phase is highest in the flamenco-model (Wilcox rank sum test *p* < 2.2*e* − 16; fig. 4F; supplementary table 2). This raises the question what causes these pronounced differences between the flamenco- and the germline-model. We suggest that recombination, due to the random assortment of cluster insertions located on different chromosomes, is responsible. Recombination among cluster sites will generate individuals with multiple redundant cluster insertions but also individuals with few or no cluster insertions. The TE will be active in these individuals without cluster insertions. Recombination thus leads to an inefficient silencing where on the average about 4 cluster insertions per diploid are necessary to furnish the majority of individuals with at least one cluster insertion. This is in agreement with our results. Under the germline-model, individuals carry various numbers of cluster insertions while in the flamenco-model the vast majority carries exactly two (fig. 4G). The few individuals with three (four) cluster insertions in the flamenco-model are likely due to multiple simultaneous insertions into the cluster at the same generation. To further test if recombination is responsible for the differences between the flamenco- and the germline-model we simulated an additional architecture: a single trap with a recombination rate of 4*cM*/*Mb* (i.e. flamenco-model with recombination; fig. 4A, setup 1). We found that the invasion dynamics of the flamenco-model with recombination are similar to the germline-model (fig. 4; supplementary table 2), confirming the important role of recombination. In Drosophila most germline clusters are located in heterochromatic regions which usually have a reduced recombination rate [Brennecke et al., 2007, Ellermeier et al., 2010]. We asked whether the absence of recombination in germline clusters has an influence on the invasion dynamics. Therefore, we simulated an architecture where we allow for recombination in the clusters that are distributed over the five chromosomes (i.e. germline-model with recombination; fig. 4A, setup 4). We however found that the invasion dynamics of the germline-model with recombination are very similar to the germline-model (fig. 4; supplementary table 2). Thus, in terms of invasion dynamics the absence of recombination in germline clusters does not confer a benefit to the host. We conclude that the flamenco-model, i.e. a single non-recombining cluster, is the most efficient architecture for stopping TE invasions. It allows for the quickest and most stable silencing response which also minimizes the amount of TEs accumulating during an invasion. Any form of recombination within/among traps, either by random assortment of chromosomes or cross-overs, renders the silencing less efficient. Note that we solely evaluated the influence of the cluster architecture. Differences in size and insertion bias were not considered (see Discussion).

**Figure 4:**
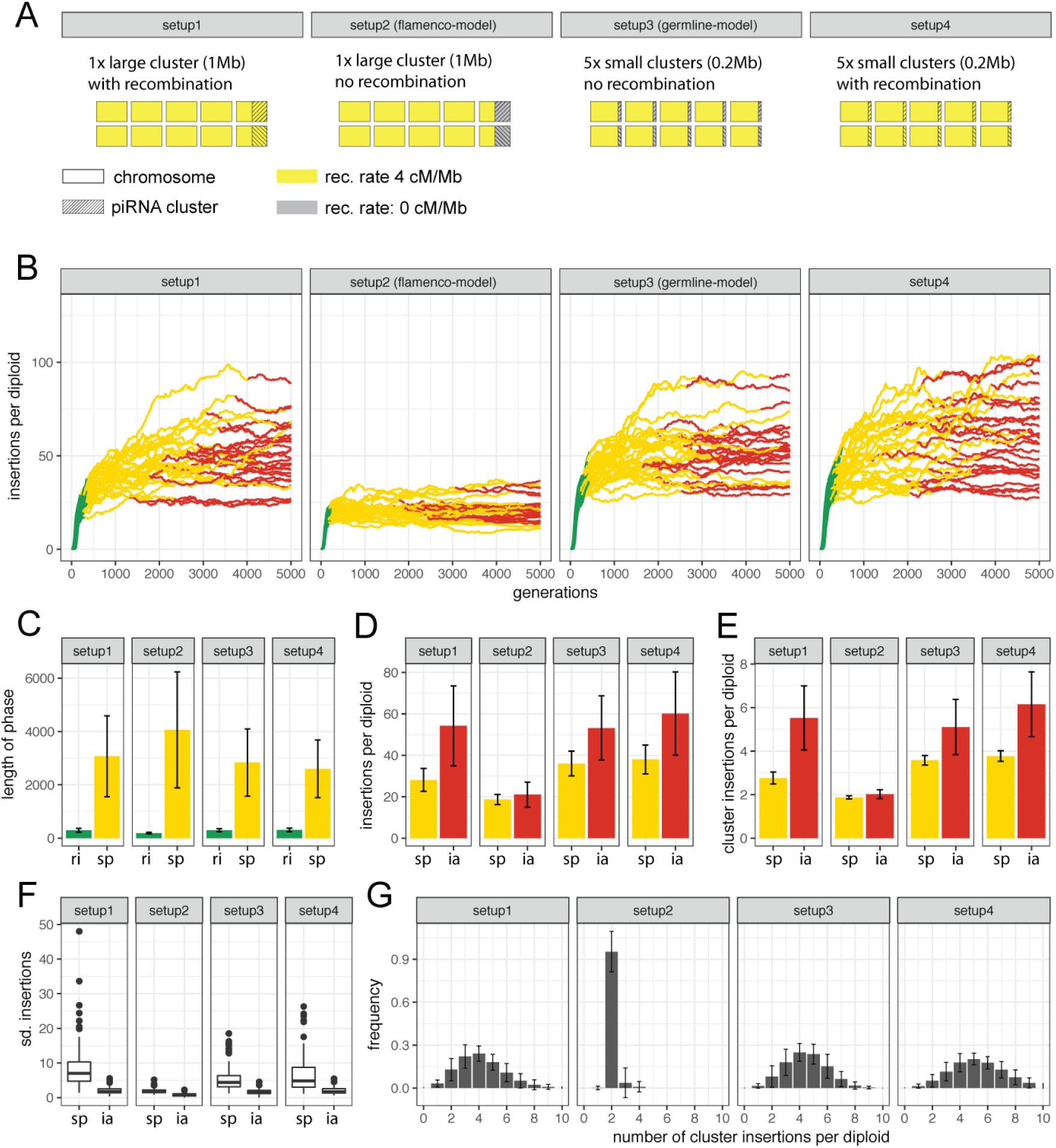
Influence of piRNA cluster architecture on TE invasions. A) Overview of the simulated architectures. Using Drosophila as example, the single non-recombining cluster resembles the architecture of the flamenco locus (setup 2) while multiple non-recombining clusters resemble the germline piRNA clusters (setup 3) B) TE abundance during invasions for the different architectures. Fifty replicates are shown. C) Length of the phases. D) TE abundance at the beginning of the phase. E) Abundance of cluster insertions at the beginning of the phase. F) Stability of the phase. G) Histogram showing the abundance of individuals with the given number of cluster insertions (at generation 1000). ri: rapid invasion; sp: shotgun phase; ia: inactive

Classic works conducted before the discovery of the piRNA pathway showed that the accumulation of TEs could be stopped by negative selection against TEs [Charlesworth and Langley, 1989, Charlesworth and Charlesworth, 1983]. In this work we show that piRNA clusters may also stop TE invasions (fig 1). It is feasible that piRNA clusters and negative selection against TEs jointly influence the dynamics of TE invasion. We therefore investigated the interaction of these two factors. Importantly negative selection against TEs could readily remove all segregating TE insertions from a population. For the following simulations we thus abrogated the previous requirement for successful invasions. Nevertheless, to avoid the stochastic early phase of invasions we initiated each simulation with 1000 randomly distributed TE insertions (frequency of insertion *f* = 1/2*N*). Initially we simulated a model where all TEs, including cluster insertions, reduce the fitness of the host by an equal amount (*w* = 1 − *xn*, where *w* is the host fitness, *x* the negative effect of TEs and *n* the TE copy number in an diploid individual). We explored the viable parameter space for TE invasions by randomly picking a negative effect (*x*) and a transposition rate (*u*). We than followed the resulting invasion up to 10.000 generations and recorded the result (fig. 5). Interestingly, in a model where solely negative selection counteracts TEs, successful invasions are only observed in a very narrow parameter space (fig. 5). If negative selection is too strong all TE insertions will be lost. If the transposition rate is too high, negative selection can not prevent the accumulation of TEs and the population will go extinct (average fitness drops to < 0.1). To extend the viable parameter space it was suggested that host fitness may not decrease linearly with TE copy numbers but exponentially instead (*w* = 1 − *xn^t^*, where *t* is an exponential factor) [Charlesworth and Charlesworth, 1983, Charlesworth and Langley, 1989, Charlesworth, 1991]. It was reasoned that ectopic recombination between TEs could have a major impact on host fitness, and that the amount of ectopic recombination may exponentially increase with TE copy numbers [Charlesworth and Charlesworth, 1983, Charlesworth and Langley, 1989, Barrón et al., 2014]. Although this exponential model extends the viable parameter space somewhat, populations still go extinct when transposition rates are high (supplementary fig. 4). Interestingly, introducing piRNA clusters into the model (with *w* = 1 − *xn*), greatly extends the parameter space over which TE invasions are feasible (fig. 5). piRNA-clusters thus prevent a rampant accumulation of TEs and rescue populations from extinction, even when negative selection against TEs is weak.

**Figure 5:**
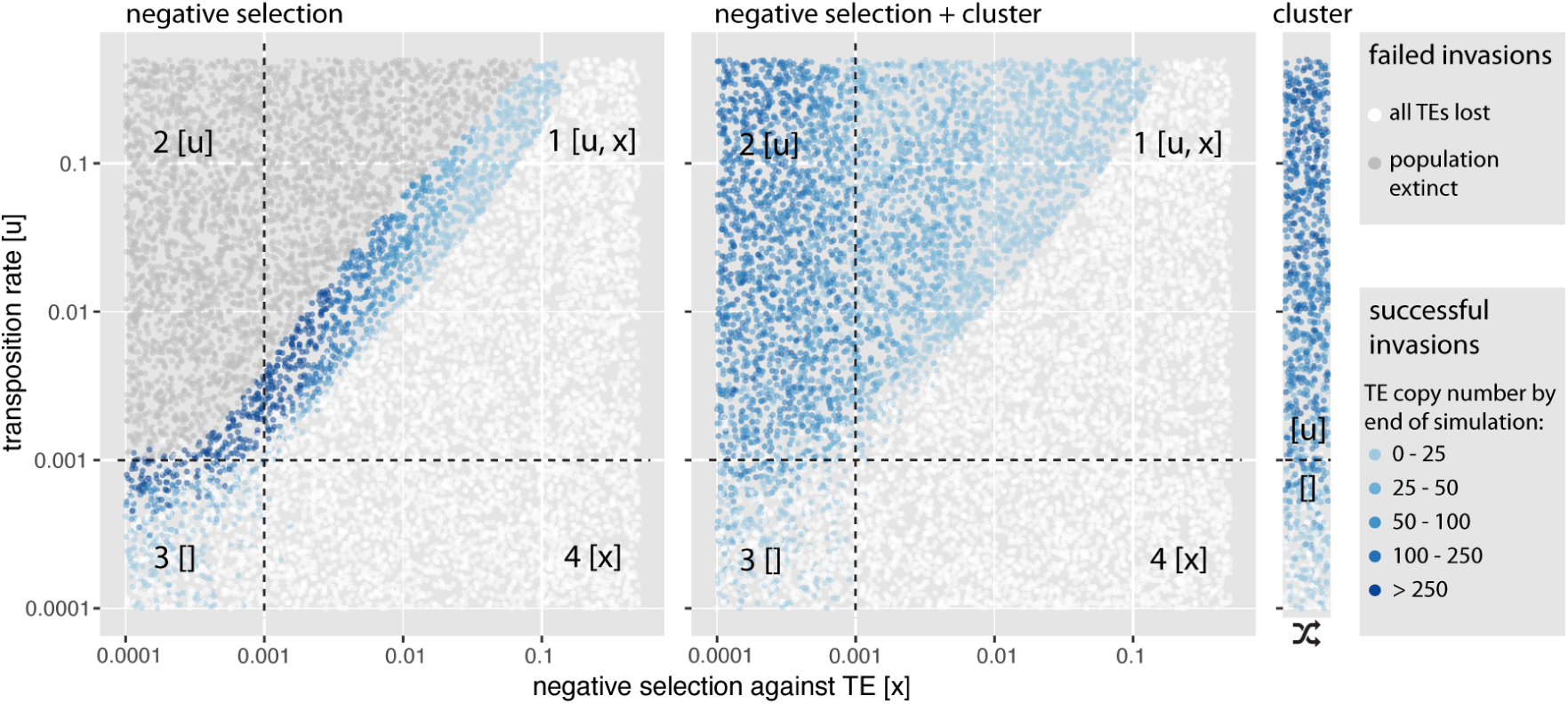
piRNA clusters protect populations from extinction due to an unchecked proliferation of deleterious TEs. Each dot represents the outcome of a single simulated TE invasion at generation 10.000. The transposition rate (*u*) and negative selection against TEs (*x*) were randomly picked. Results are shown for three different models where the following factors counteract the spread of TEs: A) negative selection against TEs B) negative selection and piRNA clusters and C) solely clusters. Dependent on the efficacy of negative selection and transposition (*N* * *u* > 1 and *N* * *x* > 1 with *N* = 1000) the parameter space can be divided into four quadrants. Factors that are effective in a given quadrant are shown in brackets. Note that piRNA clusters greatly extend the parameter space over which TE invasions are feasible.

In a model where negative selection against TEs and piRNA clusters counteract the spread of TEs three different outcomes are feasible (fig. 6A). In case negative selection against TEs is strong all TE copies are quickly purged from the population (fig. 6A, left panel). If negative selection against TEs is weak the invasion has the three phases described before (fig. 6A, right panel). Interestingly, for intermediate levels of negative selection against TEs, TE copy numbers reach a stable plateau, although fewer than 99% of individuals carry a cluster insertion. (fig. 6A, central panel). Furthermore cluster insertions are not getting fixed and the TE will thus remain persistently active. We thus found a novel equilibrium state where both piRNA clusters and negative selection against TEs, counteract the spread of the TEs. In analogy to the classic transposition-selection balance [Barrón et al., 2014] we refer to this novel equilibrium state as “transposition-selection-cluster balance” (TSC balance). Next we asked how many individuals actually carry cluster insertions during TSC balance. The fraction of individuals with cluster insertions depends on the strength of negative selection against TEs (fig. 6B; Kruskal-Wallis test at generation 10.000; χ^2^ = 520.9, *df* = 2, *p* < 2.2*e* − 16). When negative selection against TEs is strong only few individuals carry cluster insertions. Negative selection also influences the average number of TE insertions per individual, where fewer TEs are found when negative selection is strong (fig. 6B; Kruskal-Wallis test at generation 10.000, χ^2^ = 472.01, *df* = 2, *p* < 2.2*e* − 16). The allele frequency of cluster insertions is not significantly different from non-cluster insertions during TSC balance (ANOVA, effect of cluster vs. non-cluster category *p* =1; supplementary fig. 5). Finally we explored the parameter space at which TSC balance may occur (fig. 6C). Interestingly, TSC balance is mostly observed in the quadrant where both negative selection and transposition are effective (*N* * *u* > 1 and *N* * *x* > 1; fig. 6C, quadrant 1). According to basic population genetics theory a factor, such as negative selection against TEs (*x*), is only stronger than drift if the condition *N* * *x* > 1 is met [Gillespie, 2010]. This confirms that TSC balance is a three component equilibrium, where negative selection and piRNA clusters jointly counteract the proliferation of TEs. If negative selection is weak solely piRNA clusters counteract the spread of the TE (fig. 6C, quadrant 2) and if negative selection is strong all TE copies will be removed from the population (fig. 6C, quadrant 4).

**Figure 6:**
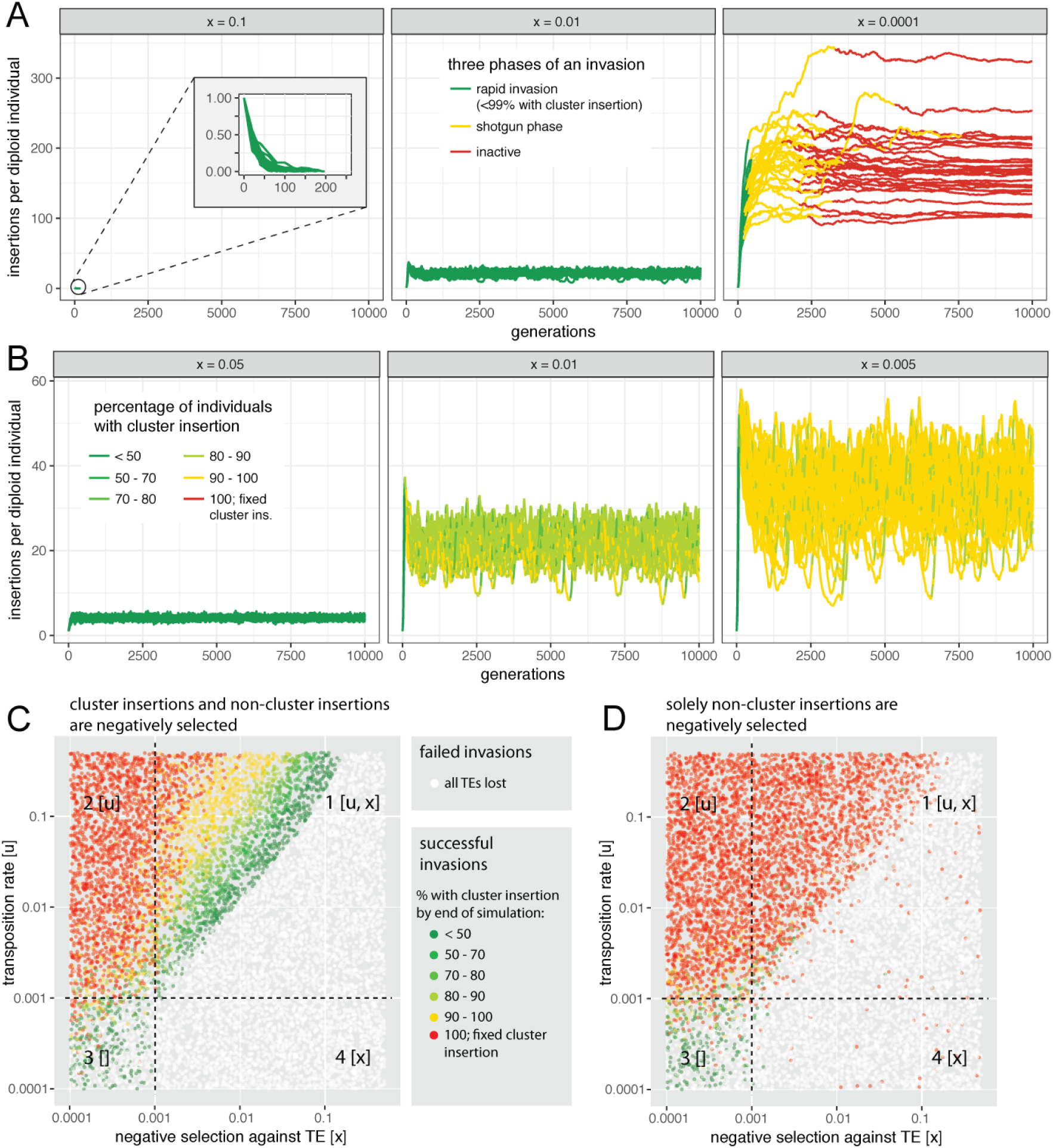
An equilibrium between <u>t</u>ransposition, negative <u>s</u>election against TEs and piRNA <u>c</u>lusters (TSC balance) prevents fixation of TE insertions. TE copy numbers may thus stabilize although only some individuals in a population carry a cluster insertion. A) Dependent on the strength of negative selection against TEs (*x*, top panel) an invasion may have three different principal outcomes: the TE may be lost (left panel), it may enter TSC balance (middle panel; these invasions never enter the shotgun phase) and the invasion may show the three typical phases (right panel). B) During TSC balance the fraction of individuals with cluster insertions depends on the strength of negative selection against TEs (*x*, top panel). Note that TSC balance prevents fixation of TE insertions, including cluster insertions and that negative selection influences TE abundance at the equilibrium. C) Parameter space at which TSC balance may be observed (green and yellow dots in first quadrant). Each dot represents the outcome of a single simulated TE invasion at generation 10.000. Dependent on the efficacy of negative selection and transposition (*N* * *u* > 1 and *N* * *x* > 1 with *N* = 1000) the parameter space can be divided into four quadrants. Factors that are effective in a given quadrant are shown in brackets. D) TSC balance is not observed when cluster insertions are neutral.

So far we assumed that negative selection is equally acting against all TE insertions, including cluster insertions. However, it is feasible that cluster insertions incur no or only weak fitness costs. In this scenario TSC balance is not observed (fig. 6D). Instead cluster insertions are quickly fixed and the TE is inactivated in most of the cases (fig. 6D). Moreover, all invasions show the three characteristic phases described before (supplementary fig. 6). Negative selection is again a major factor influencing TE abundance (supplementary fig. 6; Kruskal-Wallis test at generation 10.000; χ^2^ = 33.58, *df* = 2, *p* = 5.1*e* − 8). Interestingly the allele frequency of cluster insertions is significantly higher than the frequency of non-cluster insertions (ANOVA, effect of cluster vs. non-cluster category *p* < 2.2*e* − 16; supplementary fig. 7). In this model cluster insertions may thus have a selective advantage [see also Kelleher et al., 2018, Li et al., 2009]. Furthermore we found that most non-cluster insertions are eventually weeded out by negative selection under this model (supplementary fig. 7). Hence, mostly cluster insertions persist within populations. This model thus predicts that piRNA clusters could contain insertions from families that are not found anywhere else in the genome. In agreement with this, a careful annotation of the flamenco locus found insertions of families that are rare in *D. melanogaster* such as Pifo and Phiddipo [Zanni et al., 2013].

In summary we found that TE invasion may enter a novel equilibrium state, TSC balance, when two conditions are met: i) all TEs including cluster insertions are negatively selected and ii) both negative selection and transposition are effective in the population. During TSC balance TE copy numbers remain stable although only some individuals within a population carry a cluster insertions. Since cluster insertions are not getting fixed the TE will remain persistently active.

## Discussion

In this work we explored the dynamics of TE invasions with piRNA clusters using individual based forward simulations. We assumed that a TE is active until a member of the family jumps into a piRNA cluster, whereupon all members of the family are inactivated. This view is know as the trap model. The trap model was initially suggested by Bergman et al. [2006], even before the discovery of piRNAs, as a means to provide hosts with an adaptive immunity against TEs. Bergman et al. [2006] suggested that once a TE jumps into a cluster of nested TEs a co-suppression network is activated which silences all members of the family. One year later this hypothesis received substantial support by the discovery of piRNAs, i.e. small RNAs that mediate the transcriptional and post-transcriptional silencing of TEs [Sienski et al., 2012, Le Thomas et al., 2013, Brennecke et al., 2007, Gunawardane et al., 2007]. Based on the observations that piRNAs suppress TEs and that piRNAs are mostly produced from piRNA clusters it was suggested that a TE jumping into a piRNA cluster triggers production of piRNAs complimentary to the TE, which than silence the TE [Malone and Hannon, 2009, Zanni et al., 2013, Yamanaka et al., 2014, Goriaux et al., 2014]. This view is further supported by the finding that piRNA clusters mostly consist of TEs [Brennecke et al., 2007, Malone et al., 2009, Zanni et al., 2013]. Hence, piRNA clusters may contain the trapped remnants of past invasions. Direct support for the trap model comes from a study which found that a single P-element insertion in X-TAS (a piRNA cluster) is sufficient to silence all P-element copies in trans [Josse et al., 2007]. It is however not clear if this observation holds for all transposons and piRNA clusters. It is conceivable that for some TEs more than one cluster insertion is required to suppress activity. Small RNA biology is a dynamic research field and it can thus not be precluded that future discoveries will necessitate a modulation of the trap model.

Initially we explored invasions dynamics assuming neutral TE insertions and only later considered negatively selected TE insertions. This was done for two reasons. First, to dissect the behaviour of a complex system it is important to start with a simple model and to extend the complexity of the model only gradually by taking additional influencing factors into account [Otto and Day, 2007]. Second, the fitness effects of TE insertions remain controversial [Arkhipova, 2018]. While some studies found a negative effect of TEs [Houle and Nuzhdin, 2004, Mackay, 1989, Mackay et al., 1991, Blumenstiel et al., 2014, Yukuhiro et al., 1985] others studies obtained more ambiguous results. If TE insertions have a direct negative effect, for example by disrupting genes or promoter regions, we expect fewer TEs in the X-chromosome than in autosomes, since the negative effect of X-linked TEs is directly exposed to selection in hemizygous males. However, in Drosophila the X-chromosome has a similar TE density than autosomes, which argues against a strong direct effect of TE insertions [Kofler et al., 2012, Petrov et al., 2011]. Negative fitness effects of TEs may also arise from ectopic recombination among elements at different sites, which may lead to highly deleterious genomic rearrangements [Montgomery et al., 1987, Langley et al., 1988]. As a consequence we expect a negative correlation between the recombination rate and the TE density (assuming that rates of ectopic and meiotic recombination are correlated). While this was observed for Drosophila it was not observed for other organisms, such as Caenorhabditis and Arabidopsis [Kent et al., 2017, Quadrana et al., 2016, Laricchia et al., 2017]. This led to some doubts about the importance of ectopic recombination in containing the spread of TEs [Quadrana et al., 2016, Kent et al., 2017]. For these and other reasons Arkhipova [2018] argues that neutrality should be the null hypothesis for any evolutionary studies of TEs. Using a neutral model we found that a TE invasion consists of three distinct phases and identified factors that influence key properties of the phases. We also found that the flamenco architecture of clusters is more efficient in silencing TE invasions than the germline architecture. One testable prediction of our work is that germline-clusters should, on the average, carry more insertions per trapped family than the flamenco cluster. We also show that the population size is the major factor influencing the length of the unstable shotgun phase. Cluster insertions should thus segregate for extended periods of time in large populations. Fixation of a cluster insertion roughly requires 2 * *Ne* generations (fig. 3D). Hence, in Drosophila with an estimated population size > 10^6^, TEs that invaded recently, like many LTR families, ought to have segregating cluster insertions [Kreitman, 1983, Bowen and McDonald, 2001]. Furthermore, in case cluster insertions are segregating, the TE may still be active in the few individuals which randomly end up without any cluster insertion. We propose that this stochastic loss of cluster insertions may be an explanation for the low level of activity observed for many TE families in Drosophila (e.g. transposition rate *u* ≈ 10^−5^ [Nuzhdin, 1999]).

Later we introduced negative selection against TEs into our model and found that negative selection reduces the amount of TEs accumulating during an invasion (fig. 6B); see also Kelleher et al. [2018]). More surprisingly we found that piRNA clusters dramatically extend the parameter space over which TE invasions are feasible. piRNA clusters prevent extinction of populations from an uncontrollable proliferation of TEs. This is in agreement with the finding that piRNA clusters lower the fitness costs of TE insertions [Li et al., 2009]. In our simulations piRNA clusters account for 3% of the genome. It is feasible that smaller piRNA clusters may not be able to prevent extinction of populations over the entire parameter space. Surprisingly, we found that negative selection can have a dramatic effect on invasion dynamics. A TE invasion may enter a stable equilibrium, the TSC balance, where piRNA clusters and negative selection against TEs counteract the proliferation of a TE. TSC balance may be imagined as a form of stabilizing selection, not on a particular allele, but on the fraction of individuals with piRNA clusters. If few individuals have a cluster insertion, the TE will be highly active and novel cluster insertions will be generated. Thus the number of individuals with cluster insertions increases. If most individuals have a cluster insertion the TE will be largely inactive and negative selection will weed out TE insertions, including cluster insertions. TSC balance would be deleterious to natural populations. Because cluster insertions are thwarted from fixation, the TE will remain persistently active. Novel TE insertions will thus generate a continuous load of deleterious TE insertions in a population. This raises the question whether natural populations are actually in TSC balance for some families and how this could be identified? TSC balance predicts that only some individuals in a population will carry cluster insertions for the active TE families. This prediction could be tested by determining the abundance of cluster insertions for different families in individuals of natural populations. However, an important requirement for TSC balance is that cluster insertions are negatively selected. This could arise due to ectopic recombination between TEs or due to piRNA clusters bearing some cost to the host (e.g. metabolic cost of generating large quantities of piRNAs).

Our work highlights the profound impact of negative selection against TEs on the dynamics of TE invasions. It would thus be highly interesting to obtain reliable estimates of the distribution of fitness effects for TE insertions, ideally for cluster insertions and non-cluster insertions separately.

It has been argued that TE invasions may be stopped by hard sweeps of cluster insertions, i.e a single insertion in a piRNA cluster may be positively selected and rapidly rises in frequency [Blumenstiel, 2011, Yamanaka et al., 2014]. In this work we suggest an alternative explanation: TE invasions are initially stopped by many segregating insertions in piRNA clusters [see also Kelleher et al. [2018]]. This is in agreement with our previous work where we had the opportunity to monitor a natural P-element invasion in experimentally evolving populations of *Drosophila simulans* [Kofler et al., 2018]. The invasion plateaued around 20 generations at which time also the first P-element insertions in piRNA clusters were observed. In agreement with our model all observed cluster insertions were segregating at low frequency [Kofler et al., 2018]. However, we found cluster insertion solely for 15% of the investigated haploid genomes whereas our neutral simulations predict that two cluster insertions per haploid genome are necessary to stop an invasion. It is possible that we missed several cluster insertions due to the incomplete *D. simulans* assembly or that euchromatic P-element insertions have been converted into piRNA producing loci by paramutations [de Vanssay et al., 2012, Mohn et al., 2014, Kofler et al., 2018]. This work however raises a third possibility. The P-element invasion may have entered TSC balance. In this equilibrium state it is not expected that all individuals carry piRNA producing P-element insertions. This could be tested by sequencing the small RNAs of several individuals from a recently invaded population. Stable P-element copy numbers in the absence of piRNAs against the P-element in some individuals would support TSC balance.

We found that the number of TEs accumulating during an invasions is mostly influenced by the size and architecture of piRNA clusters. The transposition rate, the genome size, the recombination rate, the population size, and the excision rate solely had a minor influence on TE abundance. This is fortunate as it allows us to compute the expected TE abundance under the trap model for organisms with known cluster size and architecture, without having to rely on estimates for parameters that are hard to ascertain, such as the transposition rate. Comparing the expected and the observed TE abundance will allow to test whether the trap model holds for the organism of interest. *D. melanogaster* is ideally suited for this analysis as both the architecture of piRNA clusters as well as the TE abundance are known [Brennecke et al., 2007, Kofler et al., 2015b]. We first computed the expected TE abundance for germline and for somatic TEs (fig. 7A). In the simulations we assumed that germline clusters are distributed over 5 chromosomes and account for 3.5% of the genome, while the sole somatic cluster (e.g flamenco) accounts for 0.15% of the genome [assuming a flamenco size of 300kb and a genome size of 200Mb, Brennecke personal communication; [Bosco et al., 2007]]. According to these simulations germline TEs in *D. melanogaster* should have about 52 to 162 insertions per haploid genome, while somatic TEs should have about 568 to 848 insertions (90% confidence interval; fig. 7A). When comparing these expectations to the TE abundance observed in a natural population from South Africa [Kofler et al., 2015b] we found that the abundance of germline TEs fits the prediction reasonably well (fig. 7B). The TE abundance is slightly lower than expected which could be due to negative selection against TEs (simulations are based on a neutral model). However the abundance of somatic TEs is substantially lower than expected (fig. 7B). This is even a conservative estimate as our simulations did not consider that cluster insertions in flamenco need to be antisense (effectively doubling the expectations for flamenco). We thus conclude that the trap model does not account for the abundance of somatic TEs. What could be responsible for this pronounced discrepancy? Possible hypothesis are an insertion bias of somatic TEs into the flamenco locus or siRNAs contributing to TE silencing in the soma [Barckmann et al., 2018, Sultana et al., 2017].

**Figure 7:**
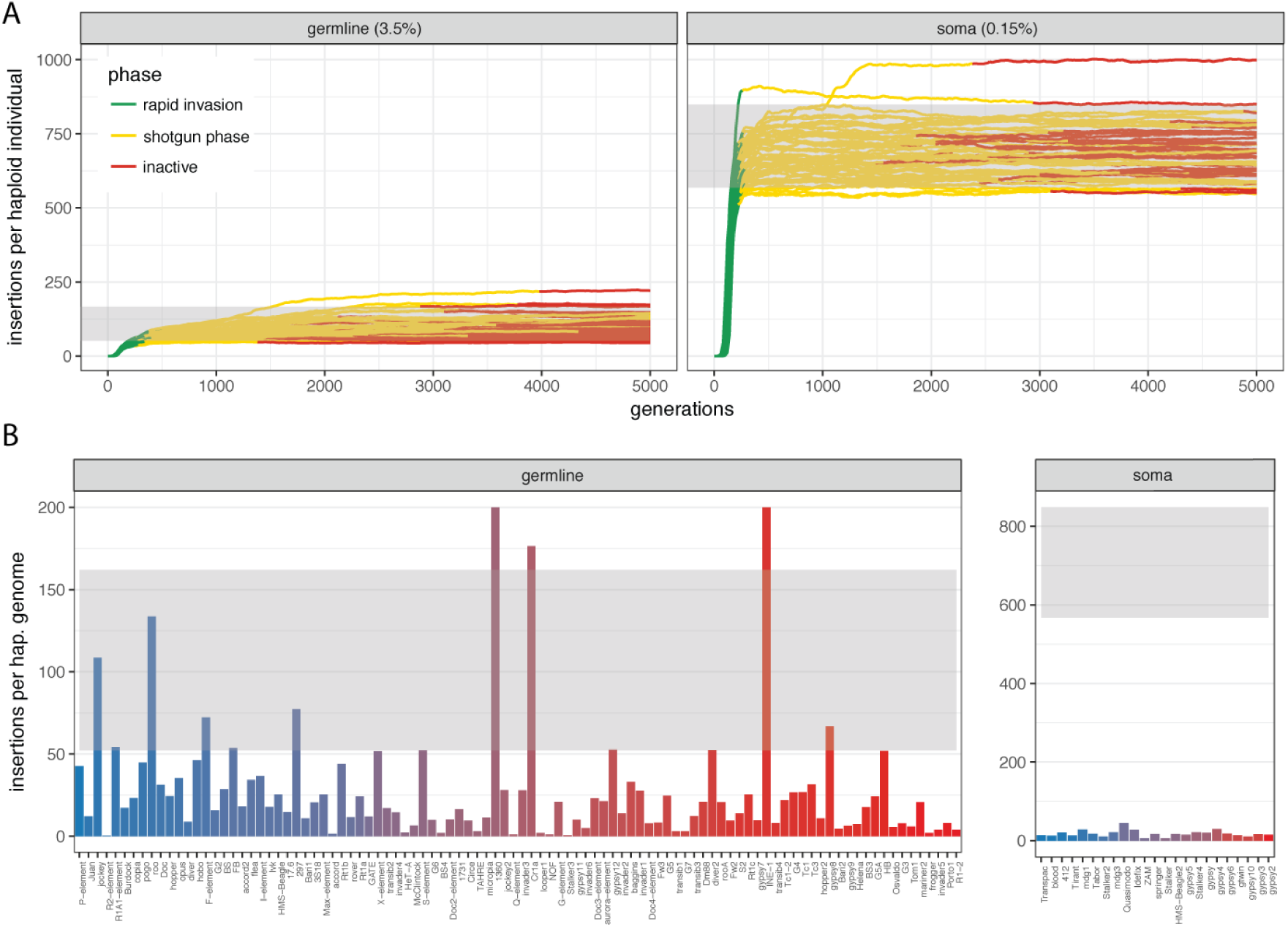
In *D. melanogaster* the trap model roughly accounts for the abundance of germline TEs but fails to explain the abundance of somatic TEs. A) Expected TE invasions for germline and somatic TEs in *D. melanogaster*. Simulated germline clusters are distributed over 5 chromosomes and account for 3.5% of the genome. A single non-recombining cluster accounting for 0.15% of the genome (flamenco) was simulated for somatic TEs. Expected TE abundance between the 5% and 95% quantile is shown in grey shade. B) Abundance of TE families in *D. melanogaster* compared to expectations. Grey shades indicate the expected TE abundance derived from the simulations (A). Color of bars indicates the average population frequency of a family (*blue* = 0.1, *red* = 1.0). Data are from Kofler et al. [2015b].

## 2 Material and Methods

### 2.1 Simulations

To simulate the dynamics of TE invasion we developed “Invade”, a novel Java tool that performs individual based forward simulations of TE invasions under different models. This tool builds on Java libraries developed for previous works [Kofler et al., 2015b, Vlachos and Kofler, 2018]. Invade allows to specify a wide range of different parameters such as the genomic architecture (number and size of chromosomes), the recombination rate, the architecture of piRNA clusters, the population size, the transposition rate, the excision rate, negative selection against TEs, and the TE abundance in the starting population. The tool also provides diverse summary statistics as output, such as the site frequency spectrum of TEs and the TE abundance in individuals of a population. At each generation Invade performs the following steps in the given order i) mate pairs are formed based on the fitness of the individuals ii) haploid gametes are generated based on the recombination map iii) TE excisions are introduced iv) novel TE insertions are introduced v) zygotes are formed vi) piRNA cluster insertions are counted vii) the fitness of the individuals is computed and vii) the output is generated [optional]. To minimize the parameter space we performed simulations with default conditions and varied solely the parameter of interest. Per default we used a genome consisting of 5 chromosomes with size 10Mb (*--genome mb:10,10,10,10,10*), a recombination rate of 4cM/Mb (*--rr cmjmb:4,4,4,4,4*), a piRNA cluster at the beginning of each chromosome with a total size of 3% of the genome (*--cluster kb:300,300,300,300,300*), a transposition rate of 0.1 (*--u 0.1*), an excision rate of zero (*--v 0.0*), neutral TE insertions (*--x 0.0*), a population size of 1000 *(--N 1000*) and 10 TE insertions randomly distributed in the starting population (*--basepop seg:10*).

### 2.2 Data analysis

Data were analyzed using custom Python scripts which are available as part of the te-tools package (https://sourceforge.net/projects/te-tools/; all scripts used in this work are in the folder sim3p). This package includes scripts for annotating the phases of the TE invasions (*phasing.py*) and computing summary statistics for the phases, such as the length of a phase and the TE abundance at the beginning of a phase (*abundance-of-phase.py, variance-of-phases.py, cluinsabundance-of-phase.py, length-of-phases.py*). Statistical analysis was performed in R [R Core Team, 2012] and visualization was done with ggplot2 [Wickham, 2016]. To identify significant changes in the site frequency spectrum between cluster and non-cluster insertions we used an ANOVA. We compared a model that includes the site-frequency class and the generation to a model that additionally includes the category “cluster vs. non-cluster insertion” as explanatory variable.

### 2.3 Details on simulated scenarious

Differences in cluster size were simulated by scaling the size of all clusters proportionally. For example to obtain clusters that account for 30% of the genome we simulated piRNA clusters with a size of 3000kb (*--cluster kb:3000,3000,3000,3000,3000*). To simulate differences in genome size we scaled the size of each chromosome and cluster proportionally. For example, to simulate a genome of size 500Mb we used five chromosomes of size 100MB and five clusters of size 3Mb (*--genome mb:100,100,100,100,100 --cluster kb:3000,3000,3000,3000,3000*). Note that this approach maintains the genomic proportion of clusters at the default value of 3%. When evaluating the impact of excision rate we kept the net transposition rate (*u*′ = *u* − *v*; i.e. transpositions minus excisions) at the default value of *u*′ = 0.1. For example to simulate 10% excisions we used a transposition rate of *u* = 0.111111 and an excision rate of *v* = 0.0111111. With an excision rate of 0% the net transposition rate is identical to the transposition rate (*u*′ = *u*).

## 3 Availability

Invade is implemented in Java and distributed under the GPLv3 at https://sourceforge.net/projects/invade/

## 4 Acknowledgments

We thank John Wakeley, Ilse Hollinger, Florian Schwarz, Julius Brennecke and Kirsten-Andre Senti for helpful comments. We thank all members of the Institute of Population Genetics for support. This work was funded by Austrian Science Fund (https://www.fwf.ac.at/) grants P29016 and P30036 to RK.

